# Challenging Backpropagation: Evidence for Target Learning in the Neocortex

**DOI:** 10.1101/2024.04.10.588837

**Authors:** Pau Vilimelis Aceituno, Sander de Haan, Reinhard Loidl, Lhea Beumer, Benjamin F. Grewe

## Abstract

Computational neuroscience currently debates two competing hypotheses to explain hierarchical learning in the neocortex: deep learning inspired approximations of the backpropagation algorithm, where neurons adjust synapses to minimize an error, and target learning algorithms, where neurons learn by reducing the feedback needed to achieve a desired target activity. While both hypotheses have been supported by theoretical studies, there is currently no empirical test that compares them directly. Here we provide such tests in the mouse neocortex by evaluating both hypotheses against experimental data at the single cell level and at the population level. At the single cell level, we conduct in vitro experiments that clarify the relationship between algorithmic learning signals and synaptic plasticity within individual pyramidal neurons. At the population level, we analyze in vivo calcium imaging data in the lateral visual cortex. By combining in vivo and in vitro data we reveal a critical discrepancy between neocortical hierarchical learning and canonical machine learning.

## 2 Main

Learning in the cortex occurs through synaptic plasticity, where the strength of connections between neurons change based on experience. Whether learning refers to new behaviors, associations between stimuli, or simply a more efficient encoding of frequent stimuli, learning always requires solving the credit assignment problem: the brain must identify the relevant synapses and adjust their strengths accordingly. Since cortical neuronal networks contain multiple layers and complex pathways between sensory input and motor output, tracing back through the network to identify which specific synapses ought to change is difficult (**Fig. 1a**). In canonical artificial neural networks, this is addressed through backpropagation (BP), an algorithm that propagates error gradients backward through the network to adjust each weight in proportion to its contribution to the output error (Rumelhart et al., 1986). While BP has proven highly effective, its biological plausibility remains contentious (Crick, 1989; Lillicrap et al., 2020; Song et al., 2024). BP relies on several assumptions that are difficult to reconcile with the biological properties of the neocortex, including precise, layer-specific error signals, symmetric feedforward and feedback weights (Liao et al., 2016; Lillicrap et al., 2016; Whittington and Bogacz, 2019), and continuous, differentiable activity, all of which contrast with the spiking, asymmetric, and locally governed nature of cortical circuits. Modern BP-inspired frameworks have relaxed these biologically implausible assumptions of classical BP and thereby put BP forth as a hypothesis to explain learning in cortex, through mechanisms such as dendritic error propagation (Sacramento et al., 2018), equilibrium propagation (Scellier and Bengio, 2017), feedback alignment (Lillicrap et al., 2016; Nøkland, 2016), and others (Whittington and Bogacz, 2017; Akrout et al., 2019; Amit, 2019).

**Figure 1:**
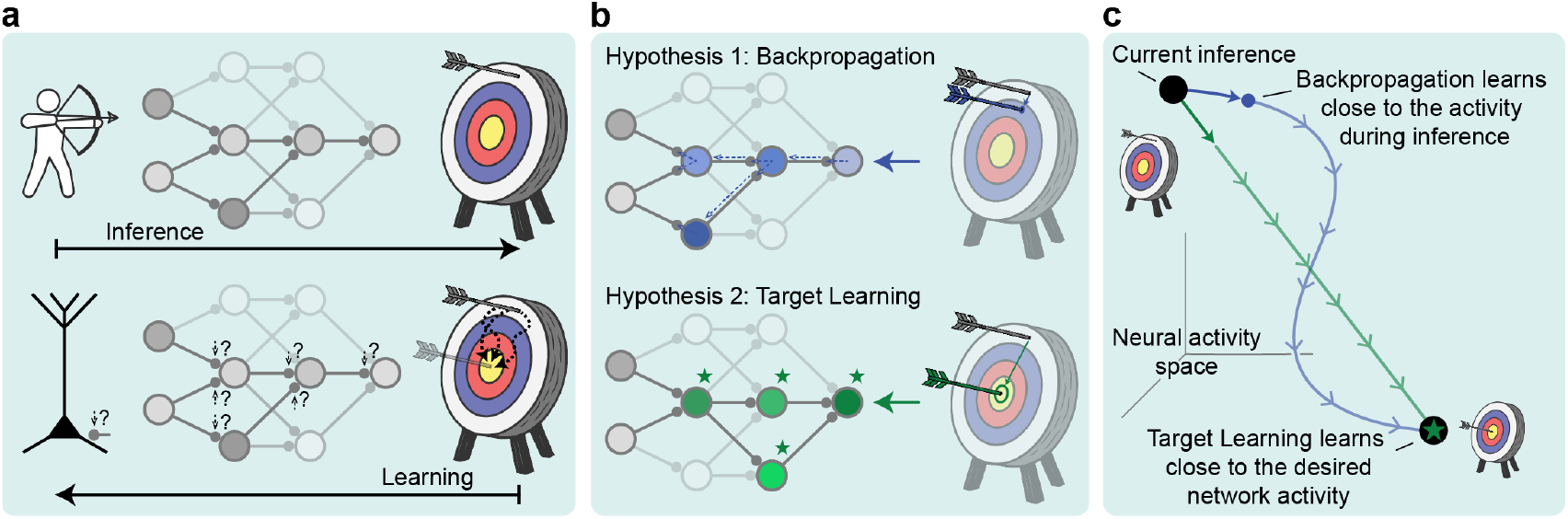
Illustration of BP and TL. (**a**) Illustration of the credit assignment problem in the cortex. (**b**) Illustration of the BP and TL hypotheses on the neural activities during learning in a network. BP relays the gradient information of the error to the individual synapses and minimally deviates from inference activity, while TL first enforces a target activity to every neuron whereafter synapses adjust to facilitate this activity. (**c**) Illustration of the BP and TL hypotheses in neural activity space. BP predicts activity during learning to be close to activity during inference, while TL predicts that activity during learning is close to the desired network activity.

In parallel, a recently growing body of work proposes an alternative cortical learning hypothesis referred to here as target learning (TL). Rather than transmitting explicit error signals, TL frameworks posit that learning occurs by imposing target activity patterns on neurons via feedback or control signals. Synaptic plasticity then adjusts the network such that these target patterns can eventually be produced without the need of an additional learning signal. Such algorithms can be traced back to data assimilation procedures (Stuart and Zygalakis, 2015), but have since reappeared in neuroscience in architectures relying on predictive coding circuits (Rao and Ballard, 1999), the FORCE (Sussillo and Abbott, 2009) and FOLLOW (Gilra and Gerstner, 2017) methods to train recurrent neural networks, and more recently the Deep Feedback Control (Meulemans et al., 2021, 2022b) and Prospective Configuration (Song et al., 2024) methods for deep networks, as well as more neuron-centric approaches (Capone et al., 2023; Saponati et al., 2025). Conceptually, TL reframes learning not as the minimization of a global error, but as the local alignment of neuronal activity with desired target states (**Fig. 1b**). Because our analysis relies on the existence of those targets, we use the term Target Learning, in line with existing experimental works (Grienberger and Magee, 2022).

Both families of algorithms have advanced our theoretical understanding of learning in neural systems (Körding and König, 2001; Richards et al., 2019; Song et al., 2024). However, different kinds of learning frameworks or circuits can approximate both BP and TL such as predictive coding (Whittington and Bogacz, 2017; Song et al., 2024), energy models (Scellier and Bengio, 2017), or dendritic error circuits (Sacramento et al., 2018; Meulemans et al., 2021), emphasizing that theory alone cannot resolve this distinction. Thus, the algorithmic principles employed by the neocortex to solve the credit assignment problem remains an open question, even after decades of debate (Crick, 1989; Lillicrap et al., 2020; Saxe et al., 2021).

In this work, we test the BP and TL hypotheses for cortical learning by examining pyramidal neuron (PN) plasticity. We first establish the mathematical differences between the two hypotheses, and derive their respective learning rules. We then test in vitro whether these learning rules align with synaptic plasticity in single PNs. Finally, we express the hypotheses in terms of observable neural activities, and test our predictions in a recent in vivo dataset.

## 3 Results

To understand how cortical neuronal networks learn, we start by denoting a mathematical formalism that captures learning rules used by single neurons. With this in mind, we detail the intracellular processes in PNs, theoretically relating teaching signals to plasticity, whereafter we conduct an in vitro experiment to directly test whether canonical assumptions of learning rules in existing BP and TL models align with PN plasticity. Next, we formulate a test that is capable of differentiating between the BP and TL hypotheses at the neuronal population level. We then select a dataset where this test can be applied in vivo, validate that this dataset contains all the necessary elements to distinguish TL and BP, and finally we perform the test.

### 3.1 Theoretical basis

To make a valid comparison between BP and TL, we first present their underlying principles at the single neuron level, and derive their respective weight update rules.

We begin by introducing some basic definitions. Both BP and TL can be understood as optimization methods that change the synaptic weights *w* of a neuronal network to ensure that the response of the network to an input sample *x* matches the desired response *y*. Mathematically, the neuronal network is represented by a function *f*(*x, w*), and for each point we can define an error term *l*(*f*(*x, w*), *y*) which quantifies how distant the response of the network is from the desired response. This error is a mathematical abstraction that might not have a direct biological correlate, making interpretation in experiments difficult (Kogo and Trengove, 2015).

The neuronal activities themselves are directly observable and require no such interpretive assumptions. To explain the BP vs TL dichotomy, three types of activity are particularly important for a given stimulus *x*: the vector of neuronal activities on the presentation of the stimulus (before learning) *a*^*S*^(*x*); the vector of target activities the neurons should ultimately produce for that stimulus *a*^⋆^(*x*); the vector of neuronal activities during learning *a*^*L*^(*x*). Now we can express both BP and TL in terms of these observable neuronal activities.

BP is a first-order gradient-descent algorithm that adjusts the weights *w* to reduce the error on each training example,

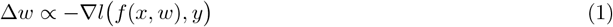

To compute this gradient, BP uses the chain rule to propagate an error signal backward through the network. At each synapse, this expresses a local learning rule of the form

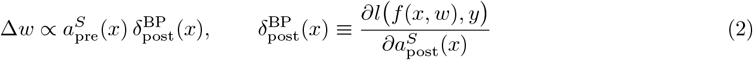

where 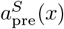 is the presynaptic activity induced by stimulus *x* before learning, and 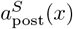 is the correspond-ing postsynaptic activity. The term 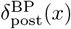 is the local error or learning signal, telling the postsynaptic neuron how much its current activity 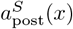 contributes to the output error and in which direction its synapses should be adjusted. This signal vanishes when the network’s output is correct, *a*^*S*^(*x*) = *a*^⋆^(*x*). Importantly, during learning in BP, the neuronal activities remain close to the activities during stimulus presentation, *a*^*L*^(*x*)≈ *a*^*S*^(*x*).

TL directly imposes the desired target neuronal activities *a*^⋆^(*x*) for a given stimulus *x* and then adjusts the synaptic weights so that the network will later produce the target activity on its own. To achieve the target activities, TL introduces a neuron-specific signal *c*(*x*) that pushes each neuron to its target activity. What TL aims to find is the signal with minimal magnitude that forces the network to the desired state,

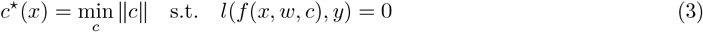

When this *c*^*⋆*^(*x*) is applied, the activities become (exactly or approximately) the target activities,

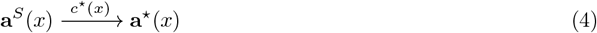

Weight updates are then performed while the network is pushed to its target activity, with the goal of minimizing the distance to this target activity state on future presentations of the same stimulus. This minimization can be done by various local learning rules (Gilra and Gerstner, 2017; Meulemans et al., 2021; Aceituno et al., 2023; Song et al., 2024), but to direct ourselves to observable neuronal activities we will use the delta rule,

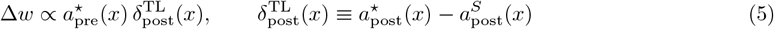

which can be derived from the mathematical principles of TL (**Supp. Text 1**). A distinctive feature of TL is that learning occurs while both presynaptic and postsynaptic activities are near their target values:**a**^*L*^(*x*)≈ **a**^⋆^(*x*). Thus, the target activity **a**^⋆^(*x*) appears explicitly in both the presynaptic and postsynaptic terms of the learning rule.

When comparing **Eq. 2** with **Eq. 5**, their similarity is apparent. In fact, the computational machinery of BP can be used to implement TL (LeCun, 1986; Ernoult et al., 2022) and vice versa (Whittington and Bogacz, 2017; Scellier and Bengio, 2017). However, doing so requires adding mechanisms and parameters to the basic algorithms, making the modified algorithms less parsimonious (**Supp. Text 2**). Therefore, any empirical evidence that favors the basic version of either BP or TL could only be supportive of the other algorithm if the extra mechanisms are also accounted for in the data. Thus, despite the similarities, the basic principle guiding how the two algorithms steer neuronal activity toward desired outcomes predicts fundamentally different observations of neuronal activity during training phases.

These differences allow for the formulation of two incompatible and testable hypotheses. For BP, neuronal activity during learning *resembles the activity during inference*, the last activity in response to a stimulus. For TL, neuronal activity during learning *resembles the activity after learning*, the target state. To evaluate these hypotheses, we will consider two approaches. First, we will investigate synaptic plasticity at the single neuron level in vitro, and infer whether this aligns with either TL or BP. Second, we will investigate neural activity at the population level in vivo.

### 3.2 Mechanisms for basal plasticity in PNs

Having established the mathematical differences between BP and TL in terms of learning rules, we now investigate how those rules relate to synaptic plasticity in individual PNs. Existing observations and theories posit that learning in the neocortex is centered on L5 PNs, which process learning signals through their apical dendrite (Spratling, 2002; Guerguiev et al., 2017; Sacramento et al., 2018; Lillicrap et al., 2020; Payeur et al., 2021; Meulemans et al., 2021).

As we have seen, the fundamental difference between BP and TL is the effect of the learning signal on the neuronal activity. While exact BP would require apical inputs that trigger plasticity but have little to no effect on the neuronal activity, TL would suggest that apical inputs alter the neuronal activity significantly (**Supp. Text. 1**). Experimental evidence shows that apical inputs strongly modulate the responses of pyramidal neurons to stimuli (Larkum et al., 1999; Williams and Holtmaat, 2019). However, these activity changes have not been directly linked to synaptic plasticity, and other mechanisms could gate plasticity (Frémaux and Gerstner, 2016).

To better understand the mechanisms by which apical inputs could affect neuronal activity and plasticity, we examine the subcellular processes that are triggered by them. At the subcellular level, apical inputs that are strong enough to elicit a local dendritic event in the form of an NMDA spike remain distant from the soma and have little effect on somatic activity (Yuste et al., 1994). However, the combination of apical inputs and a backpropagating action potential (bAP) can engage voltage-gated conductance at the apical dendrite, triggering dendritic Ca^2+^ plateau potential. These apical plateaus also elevate the somatic membrane potential, promoting bursting (BAC firing) (Larkum et al., 1999). Thus, apical inputs can determine whether a bAP will trigger a plateau potential by coincidence. Both bAPs and plateau potentials convey information about somatic activity to the synapses by propagating along the basal dendrites (Markram et al., 1997).

At the basal synapses, the depolarization induced by single bAPs or plateaus can have strong effects on plasticity. Long-lasting depolarization, as would be induced by a plateau, can effectively relieve the Mg^2+^ block of NMDA receptors during presynaptic glutamate release, thereby increasing Ca^2+^ influx into the spine (Clopath et al., 2009; Bock et al., 2022). In turn, the calcium concentration and dynamics determine long-term changes in synaptic strength (Lisman, 1989; Yang et al., 1999; Nevian and Sakmann, 2006; Graupner and Brunel, 2012). Thus, a plateau would generate enough membrane depolarization to induce long term potentiation (LTP) (Kampa et al., 2006; Clopath et al., 2009), while single bAPs fail to induce LTP in physiological conditions (Inglebert et al., 2020), and can instead induce long term depression (LTD) (Sjöström and Häusser, 2006; Chindemi et al., 2022).

To better capture the interactions between apical input, somatic bursting, and basal synaptic plasticity, we built a simplified neuron model capturing the dynamics of voltage and calcium in the apical tuft, the soma, and the synapse (**Fig. 2a, Methods 2**). Using our model we derive a learning rule similar where the postsynaptic term 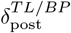 resembles that of **Eq. 5** with the target activity being represented through the ratio of bursts and single action potentials, which is itself determined by the rate of apical inputs (**Supp.Text 3**).

**Figure 2:**
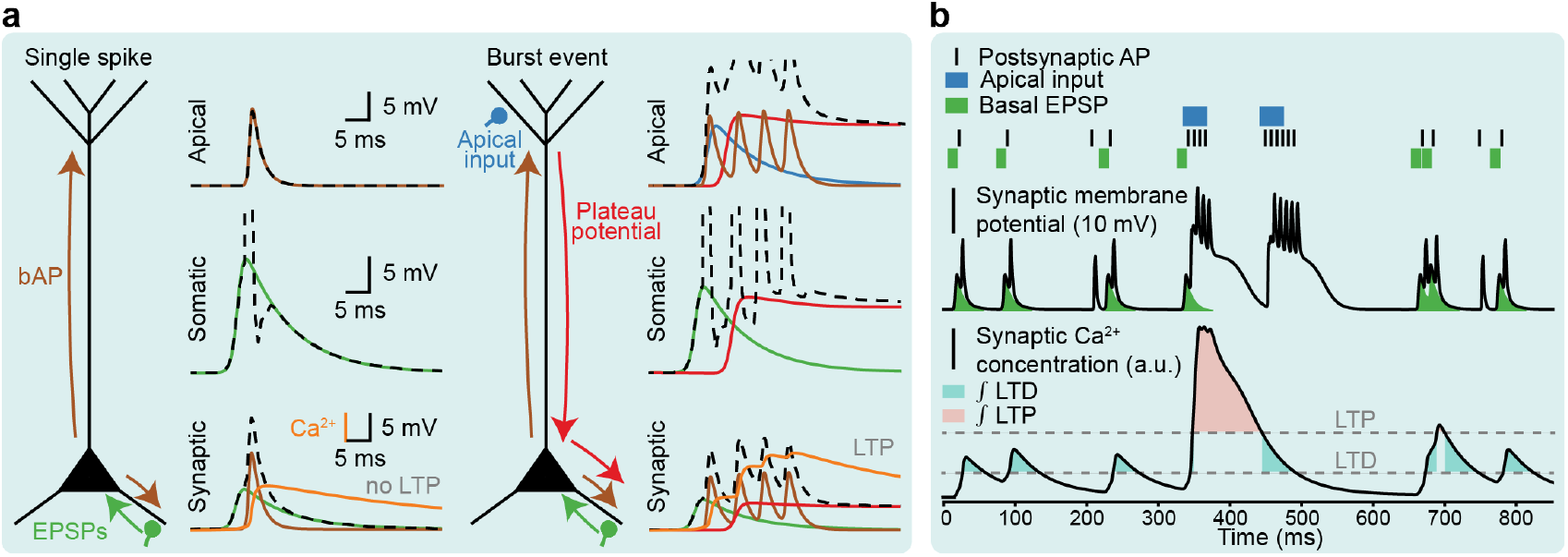
The mechanistic integration of basal and apical inputs in PNs. (**a**) Neuron schematic details synaptic depolarization events and their interactions in the PN. Responses of each compartment are shown for apical, somatic, and synaptic compartments for cases without (top) and with (bottom) top-down input to the apical dendrite. Shown for a single spike event (left) and a burst event (right) (**b**) Simulation of the effects of apical input on somatic firing rate, synaptic membrane potential, calcium influx, and plasticity. For a specific basal synapse, the causation of postsynaptic APs and apical input (top) depolarize the synaptic membrane potential (middle) and modulate the synaptic calcium concentrations leading to LTD and LTP (bottom).

Our model aligns with theoretical studies using burst-driven learning (Urbanczik and Senn, 2014; Payeur et al., 2021; Greedy et al., 2022) and apical dendritic error mechanisms (Sacramento et al., 2018; Meulemans et al., 2020; Aceituno et al., 2023) that affect both activity and plasticity. Note that the link between somatic bursts and basal plasticity is not a causal relationship, but rather a reflection of the fact that both are driven by plateau potentials (**Fig. 2b**). In the next section, we test the relationship between basal synaptic plasticity, plateau potentials, and changes in somatic activity by experimentally manipulating apical inputs.

### 3.3 Apical input directs synaptic plasticity in basal synapses

To test whether apical inputs facilitate plasticity in basal synapses and somatic activity changes experimentally, we conducted whole-cell patch clamp experiments on 12 L5 pyramidal neurons from the prefrontal cortex of six mice, and used extracellular stimulation of basal and apical afferents (**Fig. 3a, Methods 3**). Recordings were performed in physiological extracellular calcium concentrations (Inglebert et al., 2020).

**Figure 3:**
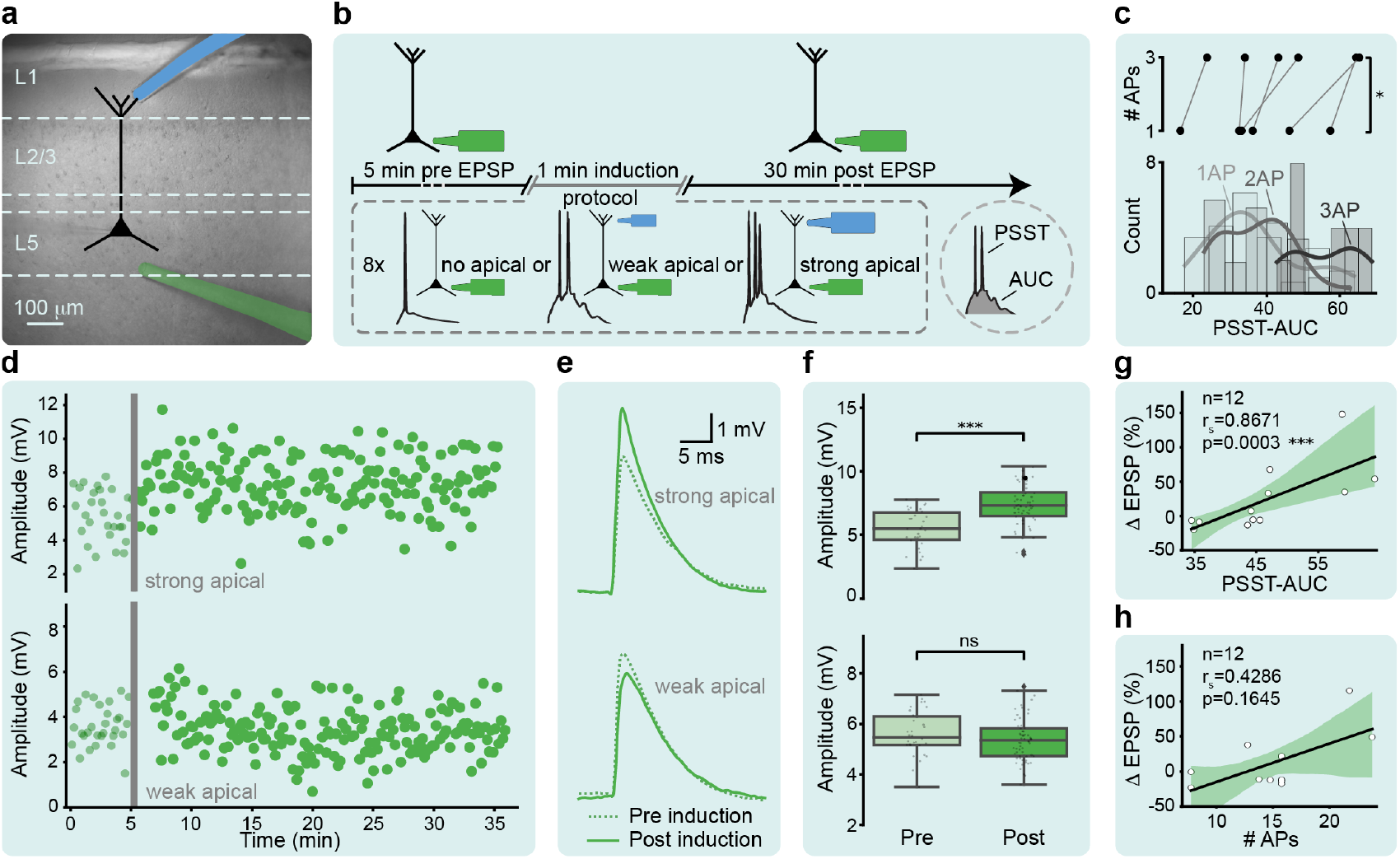
Apical inputs direct changes in synaptic strength in basal synapses and affect PN activity. (**a**) Wide-field image of mouse neocortex with an L5 pyramidal neuron sketch superimposed, and the placement of extracellular basal and apical stimulation electrodes. (**b**) Stimulation and recording protocol. (**c**) Pairwise comparison of PSST-AUC for the minimum and maximum number of somatic APs (top, Wilcoxon, *p*= 0.031). Histogram of the PSST-AUC for one, two, and three APs (bottom). (**d-e-f**) Example recordings of basal EPSP amplitudes before and after a plasticity induction event with significant increase (top) and no-significant difference in synaptic strength (bottom). (**d**) Evolution of the EPSP amplitude over time. (**e**) Mean EPSP for pre and post plasticity-induction event. (**f**) Quantification of ΔEPSP (T-test, top: *p <*0.0001, bottom: *p*= 0.229), 12 neurons from 6 mice. (**g-h**) Correlation analysis between changes in basal EPSP amplitude and PSST-AUC and number of APs, respectively (Spearman correlation, n=12, **g**: *R*_*s*_= 0.6871, *p*= 0.0003, **h**: *R*_*s*_= 0.4286, *p*= 0.1645).

We recorded baseline EPSPs for five minutes to assess initial synaptic responses. We increased basal stimulation intensity to induce one action potential and adjusted apical stimulation to trigger supra-threshold events with one to three action potentials and prolonged somatic depolarization, suggestive of a plateau potential. After eight paired stimulation-induced supra-threshold events (PSSTs), we recorded EPSPs for 30 minutes to measure long-term amplitude changes (**Fig. 3b**). We quantified somatic depolarization as a proxy for synaptic depolarization by measuring the area under the curve of somatic membrane depolarization during PSSTs (PSST-AUC).

Prolonged somatic depolarization from dendritic plateau potentials enables burst firing (Larkum et al., 1999), showing a dependency between the number of action potentials and PSST-AUC. Here, we find that the number of action potentials increases with PSST-AUC, and this change is significant even in single neurons (**Fig. 3c**, *p*= 0.031).

Applying the strong apical stimulation induction protocol facilitating burst firing increased basally-induced EPSP amplitudes indicating LTP (**Fig. 3d-f top**, *p*< 0.0001), while applying weaker apical stimulation showed no significant change (**Fig. 3d-f bottom**, *p*= 0.229). To assess basal synaptic plasticity predictors, we correlated the number of action potentials and PSST-AUC with EPSP amplitude changes (ΔEPSP). PSST-AUC showed a significant positive correlation with ΔEPSP (*R*_*Spearman*_= 0.8671, *p*= 0.0003, **Fig. 3g**), whereas the correlation with the number of action potentials was not significant (*R*_*Spearman*_= 0.4286, *p*= 0.1645, **Fig. 3h**). Thus, total somatic depolarization predicts synaptic strength changes better than the number of action potentials. In other words, the calcium plateau potential is likely the driver of plasticity with bursting as an epiphenomenon.

In contrast, we did not find clear patterns in the induced plasticity of apical synapses (**Fig. S1, Supp. Text 4**, *r*_*s*_= 0.2308, *p*= 0.4705). As we stimulate inputs to apical dendrites using extracellular electrodes, we probably jointly activate axons from the contralateral prefrontal cortex (cPFC) as well as thalamic nuclei, specifically the mediodorsal (MD) and ventromedial (VM) thalamus. Those axonal feedback projections are likely mediated by different neuron types, which could explain the heterogeneous plasticity we observed.

Our in vitro experiments show that basal synaptic plasticity in L5 PNs is driven by prolonged somatic depolarization modulated by apical inputs, consistent with modeling studies using PNs with apical inputs as learning signals (Sacramento et al., 2018; Meulemans et al., 2021). Plateau potentials and burst firing co-occur, aligning with burst-based learning rules (Payeur et al., 2021). If we interpret bursts as quantitative changes in somatic firing, we could interpret the link between AP changes and plasticity as indicative of TL. However, at the single neuron level we do not know if the changes in firing would match the target activity precisely. Furthermore, some models have proposed that even if there is a changing of firing, it could be compensated by other mechanisms (Sacramento et al., 2018; Payeur et al., 2021). We conclude that a definitive distinction between TL and BP requires observing neuronal activity in vivo.

### 3.4 Cortical reactivations isolate activity during inference and learning

To distinguish whether cortical learning aligns better with TL or BP predictions, we need experimental data that captures neuronal activity during learning, in vivo. We thus have two main requirements: (1) the data must reflect long-term neuronal changes driven by learning, and (2) the neuronal activity that drives this learning needs to be identifiable.

We adopt data from a recent experimental study which recorded somatic L2/3 PN activity from the mouse lateral visual cortex with two-photon calcium imaging (Nguyen et al., 2024). Here, mice were presented two stimuli in random order, where each stimulus lasts for two seconds followed by a gray screen for fifty-eight seconds before the next stimulus onset (**Fig. 4a**). During one session, the mouse was exposed to a total of 120 stimulus presentations, sixty for every stimulus. Importantly, the inter-stimuli intervals contain cortical reactivations: spontaneous reoccurrences of neuronal activity patterns resembling those seen during stimulus presentations (Rothschild et al., 2017), and which are known to require apical inputs (Sirota et al., 2003). These reactivations were classified according to which of the two stimuli they resembled most, and reactivations are considered local in time.

**Figure 4:**
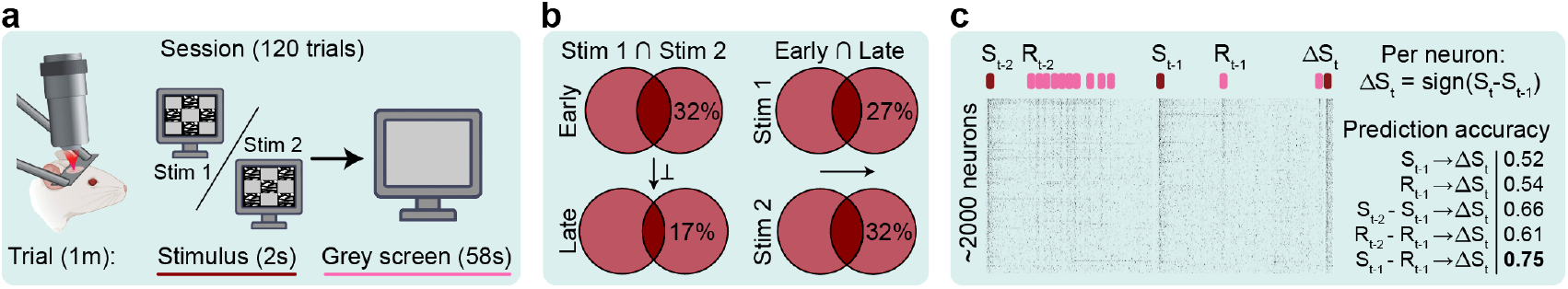
Cortical reactivations isolate activity during inference and learning. (**a**) Experimental setup from Nguyen et al. (2024) uses two alternating stimuli with resting periods, recording neuronal activity via two-photon calcium imaging to detect reactivations from stimulus-evoked responses. (**b**) The overlap between the top 5 percent of most active neurons for each stimuli decreases from early to late training (Early: 32.2 ±0.7% SEM, Late: 16.9 ±0.9% SEM) suggesting orthogonalization. (**c**) The identities of the top 5 percent most active neurons change significantly for each stimuli (Stim 1: 27.0 ±1.8% SEM, Stim 2:31.8± 4.0% SEM). (**d**) Reactivations relative to preceding stimuli best predict activity changes (pred. acc.= 0.75 vs. random = 0.5).

Note that the data captures the activity of pyramidal neurons in L2/3, not L5 as in the in vitro experiments. This is a general feature of in vivo imaging, as neural tissue is not translucid, and thus imaging deep layers is technically challenging. Although the question of which algorithm does the cortex uses is as equally valid in L2/3 as in L5, as both are part of the cortical processing hierarchy (Bastos et al., 2012), most of the modeling efforts have been focused on L5 (Larkum et al., 2001; Sacramento et al., 2018; Meulemans et al., 2021; Aceituno et al., 2023). Thus, any results obtained from this dataset need to be formulated not in terms of existing models, but in terms of general algorithmic principles implemented in abstract neuronal models (Rumelhart et al., 1986; Rao and Ballard, 1999; Song et al., 2024). With this in mind, we analyze whether this data fulfills our requirements.

For requirement (1), we assess whether the recordings capture long-term neuronal changes driven by learning. Because the in vivo dataset uses a passive exposure paradigm, one can argue that no behavioral learning is present, and thereby no learning is required. However, the study shows that the representations of the two stimuli in neuronal activity space orthogonalize during the sessions, where the representations remain stable over days, suggesting an underlying optimization objective. This orthogonalization is also visible in the overlap of highly active neurons decreasing during training (**Fig. 4b left, Methods 4.1**). This orthogonalization could arise from simple learning paradigms such as Hebbian-like learning rules (Dayan and Abbott, 2005), which would require not require sophisticated algorithms such as TL or BP. To assess whether this is the case, we note that Hebbian-like learning rules would maintain the relative ranking of neuronal activations during learning, because highly active postsynaptic neurons potentiate more (Frémaux et al., 2010). The data shows significant changes in the identity of the most active neurons over time, suggesting that changes in activity are driven by learning algorithms more complex than classical Hebbian learning (**Fig. 4b right**).

For requirement (2), the original study suggests that the reactivations are predictive of the observed long-term changes in activity, which would suggest that reactivations are the drivers of learning. This observation is consistent with previous studies showing a correlation between learning and reactivations (Chen and Wilson, 2023), as well as with causal tests in which blocking reactivation-related processes prevented animals from learning altogether (Jadhav et al., 2012), while enhancing them enhanced learning (Fernández-Ruiz et al., 2019). To verify this at the single neuron level, we tested possible predictors of activity changes in subsequent stimulus-evoked responses. Specifically, we assessed how well previous stimulus-evoked responses, reactivations, or their differences predicted the sign change of the current stimulus-evoked event, indicating the direction of learning. We train logistic regression models with every combination of events and verify them using five-fold cross-validation. We found that the difference between the previous stimulus response and its associated reactivation best predicts these changes (**Fig. 4c, Methods 4.2**), suggesting that the neuronal activity during reactivations drives learning at the single neuron level, rather than learning during stimulus presentation.

We conclude that the dataset from Nguyen et al. (2024) provides neuronal recordings capturing long-term plasticity, allowing us to isolate activity patterns during inference and learning in L2/3, which are critical for testing our algorithmic hypotheses.

### 3.5 Target learning explains reactivations in cortical networks

Drawing on our previous theoretical distinctions between BP and TL and associated hypotheses from **Section 3.1**, we can now make specific predictions about the trajectory of neuronal activity in the in vivo dataset. Since the dataset includes recordings of both stimulus-evoked neuronal responses (during inference) and cortical reactivations (during learning), we specify our predictions as follows. For BP, neuronal population activity during reactivations will resemble the most recent stimulus-evoked response for a given stimulus. For TL, neuronal population activity during reactivations will resemble the stimulus-evoked activity at the late-learning stage, when neuronal representations have stabilized.

Note that these predictions are not distinguishable during the late stages of learning, as the activity during inference is by definition very close to the activity after learning, resulting in neuronal activities during reactivations that are very similar to both previous and final stimulus evoked responses. Therefore, we focus our analysis solely on early reactivations. Every step of our data analysis is detailed in **Methods 5**. To test our hypotheses, we first project the stimulus-evoked responses into a low-dimensional space, reducing the amount of variability that is unrelated to our question. We define the first dimension as the vector from early to late stimulus-evoked responses, reflecting the learning progression. This learning-progression dimension reflects how neuronal responses evolve from the early initial activity states to stabilized and orthogonalized late training states, and therefore is the dimension of principle interest to study activity changes. We set subsequent dimensions as principal components orthogonal to this vector.

In the simpler case of a single dimension, we present the stimulus-evoked responses for one example mouse stimulus, and across all mice and stimuli (**Fig. 5b**). As expected, the stimulus-evoked responses start close to the early point and converge toward late responses. To illustrate the predictions of both algorithms, we draw theoretical lines that represent the hypothetical reactivations that would emerge from BP and TL, with the former being close to the stimulus-evoked responses at each time, and the later being fixed at the stimulus-evoked responses after learning. We find that the reactivations are closer to the TL than BP hypotheses, and stable, in line with Nguyen et al. (2024). Furthermore, we observe no learning when reactivations are not present (**Fig. 5b top**), further validating our data requirements by demonstrating the absence of learning when no reactivations are present.

**Figure 5:**
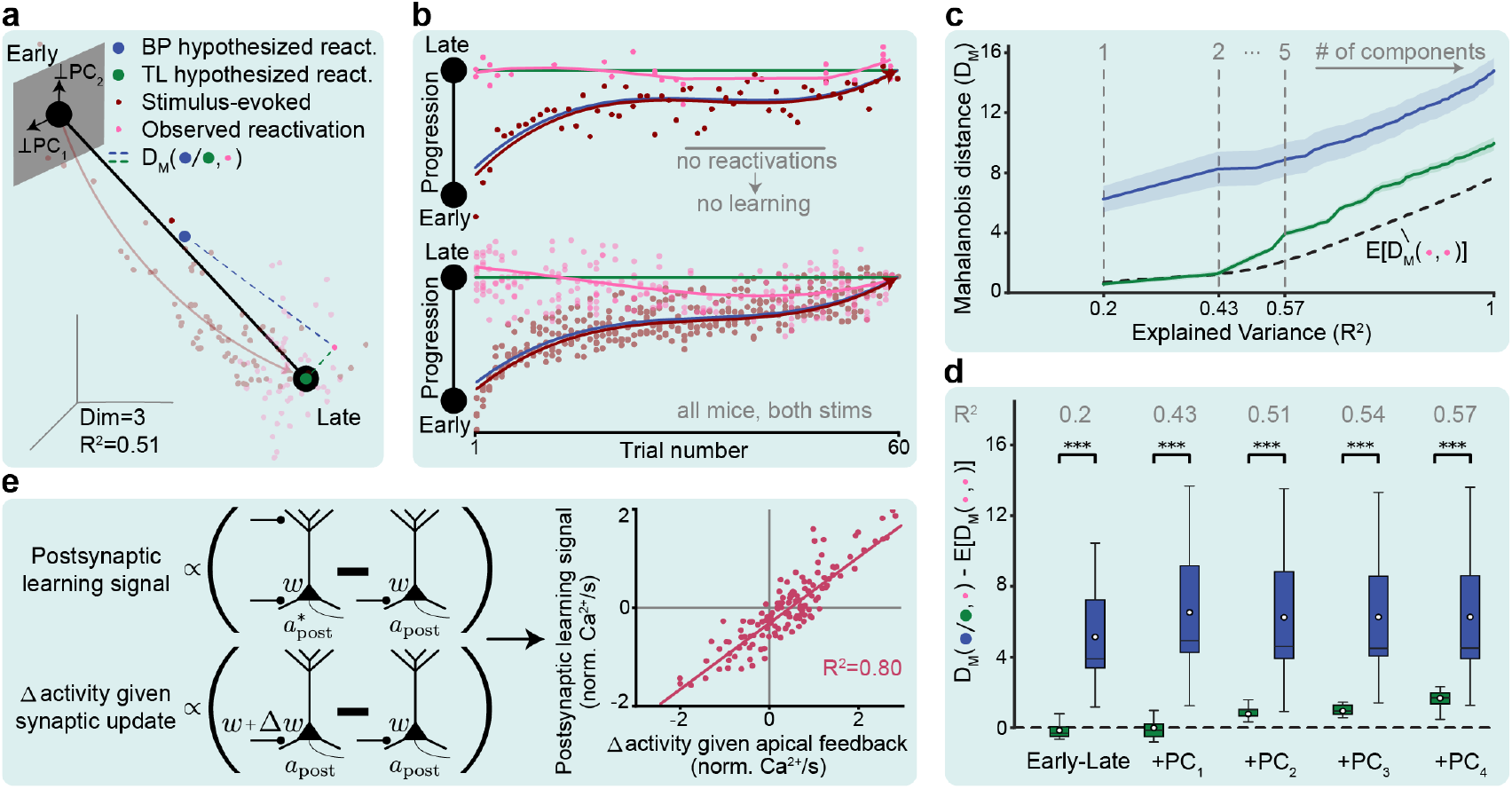
Target learning aligns with in vivo cortical learning dynamics. (**a**) Stimulus-evoked responses and reactivations trace trajectories in PC space, with the first component reflecting early-to-late changes. Distances are measured between hypothesized and observed reactivations with the Mahalanobis distance, thereby accounting for intrinsic noise present in every dimension. (**b**) Reactivations align with TL along the early-to-late component for one mouse (top) and all mice (bottom), with no learning observed without reactivations. (**c**) Mahalanobis distance increases over the number of principles components added to the dimension of measurement, the intrinsic expected distance between reactivations should be subtracted.(**d**) Mahalanobis distance shows TL explains the observed reactivations better than BP after subtracting expected distances as the number of PCs is increased, capturing a larger fraction of variance in the reactivations, (first five shown, Wilcoxon, *p <*0.0001). (**e**) Schema of the variables defining the postsynaptic learning signal (left, up) and the postsynaptic change in activity due to plasticity (left, down), and the correlation between both variables as measured in the in vivo dataset (right, *R*^2^= 0.80).

To make the comparison more precise, we next assess the distance between the hypothesized and observed reactivations, and test which hypothesis better fits the observed reactivations as we add more dimensions. We quantify the difference between the hypothesized reactivations and the real reactivations using the Mahalanobis distance (intuitively, the z-score in high dimensions) which measures the separation between data points while accounting for the inverse covariance in the neuronal activity, enhancing robustness against noisy and uninformative dimensions in the reactivation patterns. We add principal components incrementally to account for more variance of the neuronal data beyond our principal learning-progression dimension, and observe that TL consistently has a lower distance than BP (**Fig. 5d**), indicating that the TL-predicted reactivations explain the observed reactivations better. To account for the intrinsic increase in variability when considering more and more dimensions, we subtract the minimum expected distance in the Mahalanobis distance space due to the variance in the reactivations (black dotted line) from the distances measured by the two hypotheses. Within this corrected space, we observe that the TL hypothesis results in a Mahalanobis distance that is within the range of the expected distance in the space of the first two dimensions (covering a fraction of variance in the data of *R*^2^= 0.2, and *R*^2^= 0.43 respectively), and remains smaller than for the BP hypothesis for the next components. The TL hypothesis is thus much closer to the minimum expected distance (Wilcoxon test, *p*< 0.0001 for all values of captured variance). These observations are consistent across mice and stimuli (**Fig. S4**).

To further investigate how reactivations implement TL, we isolated the TL learning signals in dataset corresponding to the term 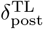 in **Eq. 5**. As our in vivo analysis suggests that reactivations represent the target (**Fig. 5e left**), we posit that 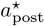 corresponds to the neuronal activity during reactivations, and 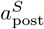 to the stimulus evoked responses. Thus, we expect that changes in somatic activity during reactivations correlate with the changes in somatic activity induced by plasticity. To test this, we calculate the difference of individual PN activities between two subsequent stimulus presentations, the change induced by plasticity, and correlate that value with the difference between the stimulus-evoked responses and the reactivations, representing the target-direction of change. We find that the two activity differences correlate very strongly and explain almost all of the variance contained in the dataset (**Fig. 5e**, *R*^2^= 0.80). This result would aligns with the predictions of our L5 neuron model in **Fig. 2b** and suggest a learning rule where the term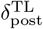 is close to that of **Eq. 5**.

## 4 Discussion

Our study combines theory as well as in vitro and in vivo experimental data, suggesting that cortical dynamics during learning align more closely with target learning (TL) rather than backpropagation (BP). This finding addresses a longstanding debate in neuroscience and machine learning about the type of learning algorithm the cortex employs (Grossberg, 1987; Crick, 1989; Körding and König, 2001; Richards et al., 2019; Lillicrap et al., 2020; Song et al., 2024), and offers a new angle to study cortical learning algorithms through reactivations. Furthermore, our in vitro experiments show that the plasticity of L5 PNs can be modulated by apical inputs and expressed through the presence of somatic bursting, complementing existing bio-plausible models of hierarchical learning (Sacramento et al., 2018; Meulemans et al., 2021; Payeur et al., 2021; Aceituno et al., 2023) and providing a biophysical mechanism for the plasticity rules that they use.

However, our study also highlights a disconnect between the modeling studies that have focused on L5 PNs, where there is a rich electrophysiology literature, and population-level data that is needed to test algorithmic hypothesis in vivo, which focuses on L2/3. Although existing in vitro studies suggest that L2/3 PN undergo similar plasticity as we found in L5 (Williams and Stuart, 2003), and imaging studies show that L2/3 and L5 PNs follow similar changes over learning in both apical and somatic dendrites (Gillon et al., 2023), future work is needed to confirm this similarities.

Our results complement previous theoretical studies which have proposed various circuits and potential architectures for learning in the neocortex, such as predictive coding (Rao and Ballard, 1999), equilibrium propagation (Scellier and Bengio, 2017), learning through dendritic errors (Sacramento et al., 2018), and deep feedback control (Meulemans et al., 2021). Importantly, we do not test the architectures or circuits themselves, but we focus instead on comparing the dynamics of PNs and cortical networks during learning. This paradigm shift is crucial, because BP and TL are mathematical frameworks which are not tied to any specific formalism, and can thus be implemented by similar network architectures or circuits (Scellier and Bengio, 2017; Whittington and Bogacz, 2017, Whittington and Bogacz, 2019; Song et al., 2024). By considering only the neuronal activity patterns of TL and BP, we argue that our study is able to clarify the algorithmic principles driving learning in the neocortex.

However, by focusing solely on PNs, we abstracted away the specific circuit requirements needed to implement this family of algorithms. To address this gap, future work shall clarify the circuit-level role, origin, and generation of feedback signals in TL architectures (Rao and Ballard, 1999; Scellier and Bengio, 2017; Meulemans et al., 2022b). We provide a first step in doing so by measuring apical plasticity in L5 PNs, which has been hypothesized to mediate feedback signals (Williams and Holtmaat, 2019; Meulemans et al., 2021; Aceituno et al., 2023). However, we did not find any consistent relation between PN activity and apical plasticity, most likely due to the unspecific nature of our stimulation. Future experimental research needs to disentangle the contributions of different afferent feedback projections to elicit apical inputs and drive plasticity, for example, using projection-specific optogenetic methods.

Finally, we highlight that in contrast to previous theoretical studies (Whittington and Bogacz, 2017; Scellier and Bengio, 2017; Sacramento et al., 2018; Meulemans et al., 2021; Song et al., 2024), the in vivo task that we consider here is unsupervised and simple, as in classical models (Cottrell et al., 1987; Rao and Ballard, 1999). More complex tasks most likely would require more complex targets, making such tests more convoluted. For example, in a decision task with two stimuli with a specific movement associated to each stimulus, the final target activity might not be clear from the beginning, as the animal needs a neuronal activity for each stimulus, a neuronal activity for each movement, and eventually a single target for each contingency. Thus, we might have target activity that changes as the animal learns the task. This would require a more sophisticated analysis, and possibly in vivo data collected from multiple areas. Importantly, there are studies that could provide the necessary data if we focus on spontaneous activity.

In summary, our study combines deep learning theory with in vivo cortical data and in vitro electrophysiology. Our investigation bridges multiple scales of neuronal learning suggesting that cortical hierarchical networks learn by enforcing targets rather than correcting errors, thus differing from their canonical artificial counterparts.

## 5 Author contributions

P.V.A. conceptualized the project and performed the mathematical analysis. P.V.A. and S.d.H. developed the in vivo data analysis. S.d.H. developed the neuron model and performed the data analysis. R.L. conceptualized and performed the electrophysiological experiments, and provided biology insights. L.B. aided the in vivo data analysis. S.d.H. created the figures. B.F.G. supervised the project. P.V.A., S.d.H., R.L., and B.F.G. wrote the paper.

## 6 Author information

The authors have declared no competing interest.

## 7 Data and code availability statement

Experimental data and analysis code will be made public upon acceptance

## 8 Acknowledgements

This work was supported by the Swiss National Science Foundation (B.F.G. CRSII5-173721 and 315230 189251, P.V.A. 182539funding), ETH project funding (B.F.G. ETH-20 19-01), and a Forschungskredit from the University of Zürich (P.V.A. FK-24-122).

## Methods

### 1 Testing alignment between reactivations and target learning rules

TL learning rules often follow an equation of the form

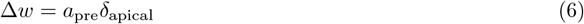

where *δ*_apical_ is an apical input, which is similar in form to **Eq. 5**. In contrast to the algorithmic rules from **Eq. 2** and **Eq. 5**, the apical input usually has an effect on the postsynaptic activity, which in the experimental literature is modulatory (Larkum et al., 1999), but is often formalized as additive in computational studies (Sacramento et al., 2018; Meulemans et al., 2021), giving

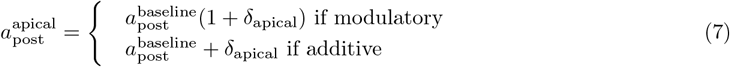

where 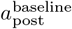 the postsynaptic activity at baseline apical input. In the in vivo data we do not measure apical inputs directly, so we must use the somatic activity only. To isolate the apical input, we can rewrite the previous equation as

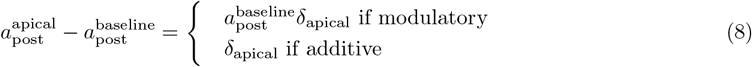

Now we have isolated the change in activity due to the apical inputs, we can isolate the apical inputs for two subsequent presentations of the same stimulus with plasticity in-between. Assuming that the activity in response to a stimulus is done at baseline activity,

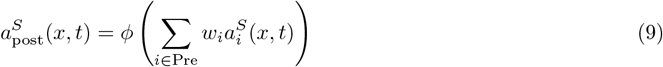

where *t* is the stimulus count, *ϕ* is the activation function of the neuron, and Pre is the set of presynaptic neurons (note that in this section we use a slightly different notation than in the rest of the document to avoid confusing stimulus identity with presentation). Assuming a similar activity of the presynaptic neurons, we can isolate the *δ*_post_ term in Eq. 6 as

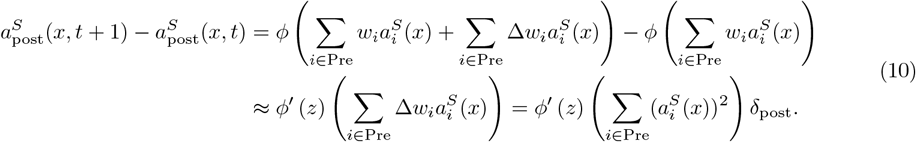

Under the assumption that the activity during stimulus presentation is at baseline apical input, we can then combine **Eq. 10** and **Eq. 8** and recover

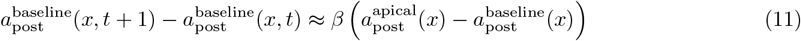

where the term *β* depends on unknown factors such as the activation function, the presynaptic activities, and the modulatory/additive nature of the apical input. In any case, we do not need to identify the form of *β*, only show that it is positive which we can do by correlating the left and right terms. To do so in our dataset, we need to assume that 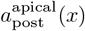 is the activtiy during reactivations.

### 2 Modeling basal synaptic plasticity and pyramidal neuron dynamics

We model basal synaptic plasticity in pyramidal neurons (PNs) based on synaptic calcium as a key driver, adopting a plasticity function *P* from Graupner and Brunel (2012) that integrates calcium dynamics to determine LTP or LTD. The synaptic strength change is computed as

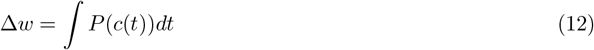

where ≈*w* is the synaptic strength change and *c*(*t*) is the calcium concentration.

Synaptic calcium influx is modeled via NMDA receptor (NMDAR) dynamics, regulated by presynaptic glutamate release and postsynaptic depolarization, using

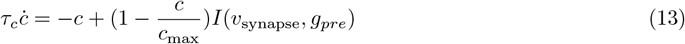

where *τ*_*c*_ is the calcium time constant, *c*_max_ is the maximum calcium concentration, and *I* represents NMDAR-mediated influx (McRory et al., 2001; Slutsky et al., 2004), with *g*_pre_ as glutamate presence and *v*_synapse_ as plateau depolarization.

Postsynaptic membrane depolarization is modeled across synaptic, somatic, and apical compartments, incorporating EPSPs, backpropagating action potentials (bAPs), and plateau potentials via

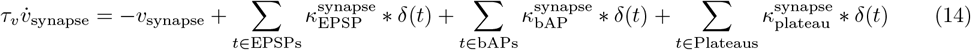

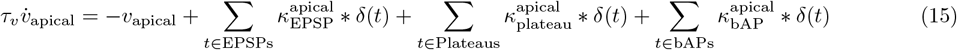

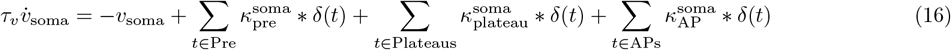

where *τ*_*v*_ is the membrane time constant, *δ*(*t*) is the Dirac delta, and *κ* denotes compartment-specific responses (see **Table 1** for hyperparameters).

**Table 1.**
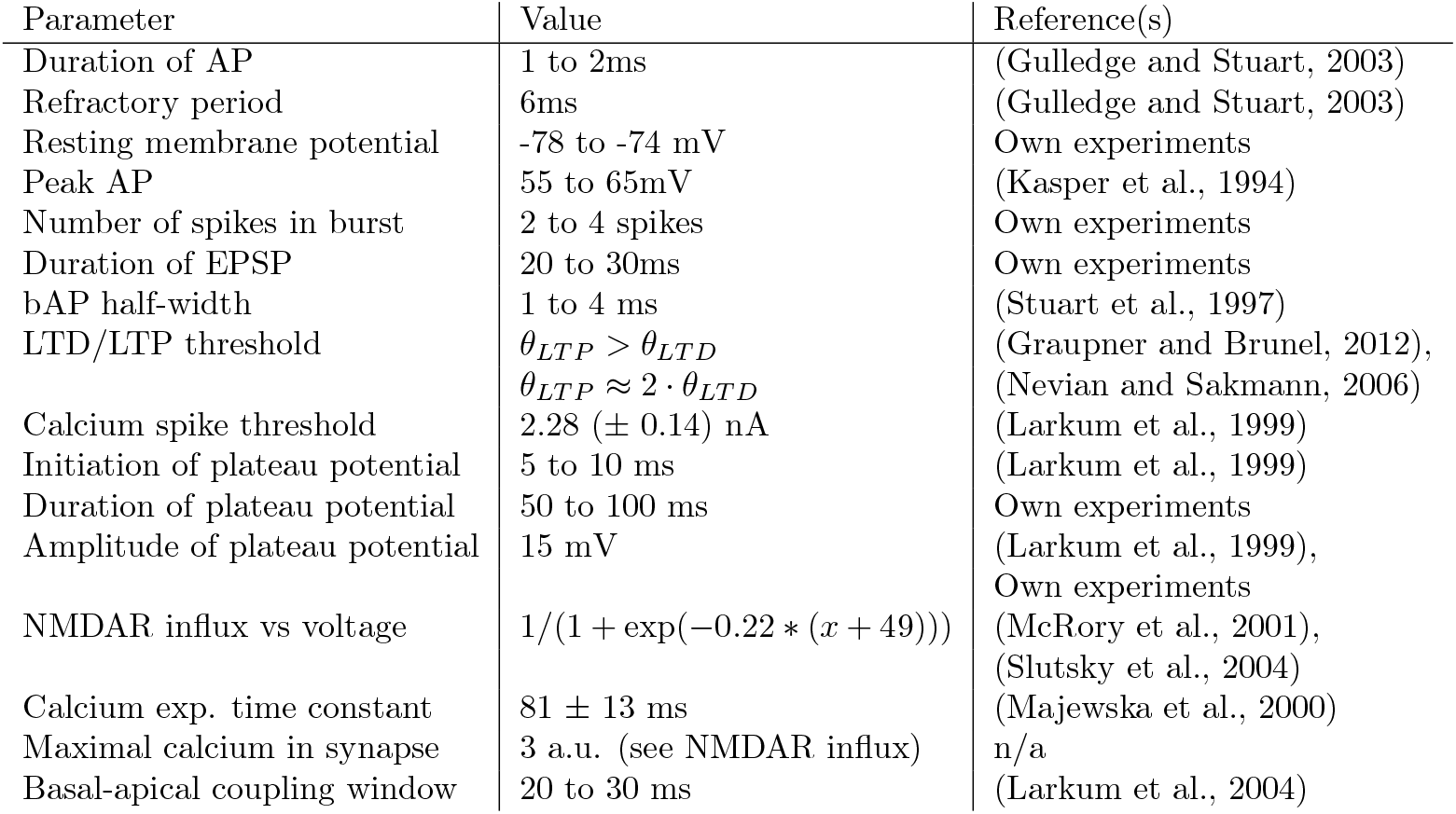
Hyperparameters of the neuron model with their associated value and references.

Plateau potentials, triggered by coincident bAPs and apical inputs via voltage-gated calcium channels (VGCCs), propagate depolarization, facilitating bAP-activated Ca^2+^ spike (BAC) firing. This bidirectional coupling between compartments modulates synaptic plasticity through NMDAR-regulated calcium influx, with apical input enhancing firing rates and plasticity when paired with basal inputs, consistent with Larkum et al. (2004).

### 3 Electrophysiology

Male B6/J-Crl1 mice of strain C57BL/6J (orig. Charles River) were bred and maintained at our in-house facility. Animals were kept in their home cages under normal day/night conditions (light: 7am –7pm). Mouse experiments were approved by the institutional animal care and use committee of Anonymous Institution. All experiments were conducted with the approval of the Anonymous State Veterinary Office.

Mice (11-13 weeks old) were deeply anesthetized using Isoflurane and decapitated. The brains were dissected from the skull and submerged in ice-cold high-sucrose artificial cerebrospinal fluid (ACSF) during acute slice preparation. High sucrose ACSF containing (in mM): 75 Sucrose, 10 Glucose, 80 NaCl, 26 NaHCO3, 7 MgSO4, 2.5 KCl, 1 NaH2PO4, 0.5 CaCl2, fumigated with oxycarbon (95% O2, 5% CO2). Coronal slices (300 µm) were drawn from PFC (+1.4 to +2.6 from bregma) using a VT1200 S vibratome (Leica) and incubated for 30 minutes in oxygenated holding ACSF at 36°C. Holding ACSF containing (in mM): 15 Glucose, 120 NaCl, 26 NaHCO3, 0.9 MgSO4, 2.5 KCl, 1.25 NaH2PO4, 1.3 CaCl2, fumigated with oxycarbon (95% O2, 5% CO2). Magnesium/Calcium ratio was chosen according to Inglebert and Debanne (2021). Slices rested for at least 45 minutes in holding ACSF at room temperature prior to electrophysiological recordings.

During recordings, brain slices were continuously perfused with oxygenated holding ACSF at 36°C. Neurons were visualized with differential interference contrast (DIC) using an Olympus BX61 WI microscope with an Olympus 60x water immersion objective and a CellCam Kikker MT100 (Cairn) DCIM camera. L5 pyramidal neurons (PN) in the PFC were identified based on morphology and distance to the pia and patched with glass pipettes with resistances of 4.8-6.4 MΩ filled with intracellular solution containing (in mM): 10 HEPES, 20 KCl, 117 K-gluconate, 4 Mg-ATP, 0.3 GTP, 10 Na-P-creatine. For morphology reconstruction (**Supp. Fig. S2**), the intracellular solution also contained 0.2% Biocytin. For Ca^2+^ imaging experiments, the intracellular solution also contained 200 µM Oregon Green™ 488 BAPTA-5N. Whole-cell, somatic patch-clamp recordings were acquired in current-clamp mode using an Axon MultiClamp 700A amplifier and digitized using an Axon DigiData 1550 B digital converter. Data was sampled at 50 kHz and filtered at 3 kHz. Typical resting membrane potentials were between −64mV and −72mV, and access resistance ranged from 15 MΩ to 25 MΩ. Recordings in which neurons depolarized > 10% from resting potential or access resistance changed > 15% from initial value were discarded. Bridge potentials were compensated, and liquid-junction potentials were not corrected for. Depolarizing and hyperpolarizing currents were injected to characterize neurons further. Only neurons showing L5 PN-typical *I*_*h*_-currents were considered.Theta-barrel glass-pipettes with openings of 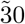 µm were placed at 50 µm – 80 µm from the patched neuron to stimulate afferents targeting basal dendrites as well as in cortical layer 1 to stimulate apical afferents with synapses onto apical tuft dendrites.

After stabilization of resting membrane potential, apical and basal stimulation strength was adjusted to elicit EPSPs with amplitudes in the range of 2-3 mV. This usually required 100 µs voltage pulses in the range of −1 to −3 V followed by immediate 100 µs voltage pulses of +1 to +3 V for basal stimulations. Apical stimulation had the same shape but ranged from −6 to −8 V and +6 to +8 V, respectively. Next, the required strength of basal stimulations to induce a single action potential was determined. For this, tripling basal stimulation amplitudes and stimulating 3 times at 100 Hz would sometimes elicited a single action potential. Due to focal stimulation (theta glass pipettes with small opening), extracellular electrical stimulation was often not sufficient to trigger action potentials. In these cases, additional brief current injections into the soma via the patch pipette were used to trigger an AP. Finally, apical stimulation strength was increased until multiple action potentials or plateau potentials were induced.

After all required stimulation amplitudes were set, baseline EPSP amplitudes were recorded for 5 minutes with alternating basal and apical stimulations every 5 seconds followed by 8 combined, strong basal and apical stimulations with a 10 second interval, each eliciting a plasticity induction event. Next, EPSPs were recorded for at least 30 minutes (same as baseline recordings).

Analysis of electrophysiology traces was performed using individual Python scripts and the pyabf library. The correlation analyses were performed with the seaborn library (regplot) and the statistical analysis with the scipy library (spearmanr).

Analysis of PSST-AUC with different numbers of action potentials was performed on a subset of neurons in which PSSTs occurred with different amounts of APs (1vs2, 1vs3, or 2vs3). PSST-AUCs were averaged for events with min. and max. number of APs respectively and plotted as 2 data points per cell.

In the case of cell morphology reconstruction (only performed in 8 of 12 neurons), slices were kept in 4% PFA for 24-32 hours and stained overnight with Streptavidin, Alexa Fluor 350 conjugate (invitrogen) and mounted on commercially available microscopy slides.

Calcium imaging was performed in a subset of neurons (n = 6 neurons, 4 mice) after post-induction EPSP recordings where completed by imaging pipette-filled OGB5-N using a Lumencor SPECTRA Light Engine (GFP/FITC 475/28 nm) for excitation and a ORCA-Lightning Digital CMOS camera (C14120-20P, Hamamatsu) for image acquisition at 50 Hz. Imaging data was pre-processed (background-subtraction, bleaching correction) using Fiji/ImageJ2. Extracted fluorescence traces were further processed and analysed using individual Python scripts. A correlation analysis between somatic calcium recordings (derived as ΔF/F AUC) and somatic membrane potential AUC showed a significant correlation (**Supp. Fig. S3**, *R*_*Spearman*_= 0.8545, *p*= 0.0016) and was used to infer calcium levels during the actual plasticity induction events using linear regression.

Group data are expressed as mean ± s.e.m. unless otherwise stated. Statistical significance was calculated using paired or unpaired t-tests as well as Spearman correlation.

### 4 In vivo population data set validation

We measure the alignment between the activity during learning and during inference or stimulus presentation in the cortical in vivo data available from Nguyen et al. (2024). We modify the existing code to fit the analysis described in the main text and keep the pre-processing as in the original publication (Nguyen et al., 2024). For all cortical data included in our analysis, we used only the top thirty percent of neurons responding to the given stimulus, more than the ten percent as in Nguyen et al. (2024). We denote the early response centroid as the average neural activity of the first three stimulus-evoked responses, and late response centroid as the average neural activity of the last three stimulus-evoked responses. We will use **r** to denote the stimulus-evoked responses and *ρ* to denote reactivations.

#### 4.1 Neural activity changes are not directly dependent on the initial neuronal activity

To assess whether credit assignment is necessary changes in neural patterns, we ranked neurons by activity of the first and last three stimulus-evoked responses per mouse, and computed the amount of overlapping neurons that were in the top 5% most active for both early and late trials. We show the average amount of overlapping neurons between early and late stimulus-evoked responses across mice in **Fig. 4b left**. To analyze the changes in stimulus representations, we also computed the overlapping top 5% of neurons in early and late trials between the two stimulus types in **Fig. 4b right**. Both overlaps are low, suggesting that simple Hebbian learning is not enough to explain the neural activity changes.

#### 4.2 Reactivations predict learning at the single neuron level

To test whether reactivations play a causal role in shaping future stim-evoked activity, we predicted the sign of change (+1 for increase, −1 for decrease) of each neuron for consecutive stimulus-evoked responses. For each day and mouse, we trained numerous logistic regression models with a variety of combinations of predictors to predict this sign of change neuron per event. We used a five-fold cross-validation strategy. For each model type, we aggregated the model accuracies across all sessions and mice. Note that across all mice and days, there were fewer reactivations than stimulus-evoked responses. For balanced comparisons, we calculated a target vector, the direction of for the next stim-evoked response relative to the current response, only for trials that contained at least one reactivation in between. In all models, therefore, the test dataset across all five folds had dimensions of (#*reactivations*× #*neurons*). In all models the target vector remained the same, whereas the set of predictors differed. For the set of predictors, we tried the following combinations: (1) Current stim-evoked response; (2) Current reactivation; (3) Current stim-evoked response minus previous stim-evoked response; (4) Current reactivation minus previous reactivation; (5) Current stim-evoked response minus current reactivation.

Granger causality was applied to assess whether reactivations enhance predictions of future stim-evoked plasticity, defined by a probability shift when reactivation data is included:

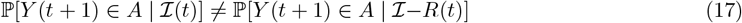

where *Y*(*t*+ 1) is the future change, ℐ (*t*) includes past stim-evoked and reactivation activity, and ℐ − *R*(*t*) excludes reactivations.

### 5 In vivo hypothesis testing

To compare the reactivations with the stimulus-evoked responses we consider them as vectors in a space where every dimension corresponds to the activity of one neuron. In this vector space, we can define both the reactivations and the activities, but the baseline activity for single neurons during rest periods – which is when the reactivations take place – is not the same as during active periods (Nguyen et al., 2024).

#### 5.1 Hypothesis testing in one dimension

The measure of progression is shown in **Fig. 5b top** using data from sample mouse ‘NN8’ for stimulus 2 from the recording date 12-03-2021. For the plots over all subjects and stimuli, we normalize the length of the line early-to-late and all points in that line. As reported by us and in the original publication (Nguyen et al., 2024), the data for other mice is looks similar (**Fig. 5b bottom**).

#### 5.2 Quantitative hypothesis comparison

We can use projection onto the early-to-late vector and PC components orthogonal to it as a dimensionality reduction procedure that maintains the space where our hypotheses live (the early-to-late vector) as well as other neural variability. In this space, we can measure the distance between the activity that would be expected in both our hypotheses and what we observe in the data.

Our approach is thus to project the neural activity of the reactivations into the space given by

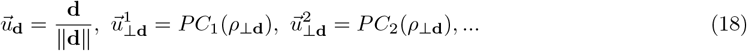

where *PC*_*i*_ is the *i*th principal component operator that is applied to the data projected on the component orthogonal do **d**. In this new basis, we can measure the noise-adjusted Euclidean (Mahalonobis) distances

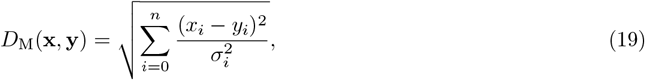

where 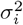 is the variance captured by the component *i*(with *i*= 0 being the early-to-late direction). We compute the distances between the reactivations and the target activity (the late stimulus response, 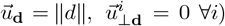, and the inference activity (as given by the stimulus-evoked responses), corresponding to the BP or TL hypotheses respectively. We can thus obtain a distance value for each reactivation for each hypotheses, and number of components (**Fig. 5c,d**) and a Wilcoxon t-test to evaluate whether their difference is significant (**Fig. 5d up**).

Since the intrinsic distances between points in a measurement with noise grow with the number of dimensions considered, the distance between our hypotheses and the reactivations is expected to grow even if one of the hypothesis is valid. To correct for this effect, we subtract the expected distance between two samples of the same point with different noises. In Mahalanobis space with normally distributed residuals and a covariance matrix normalizing variances, the squared distance between random points follows a chi-squared distribution with *n* degrees of freedom. With the covariance incorporated, the expected squared distance scales as *n*, and the expected distance (its square root) adjusts to 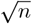. Note that there is a correction for finite dimensions, which reduces the distance to 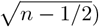, which we subtract from observed distances to isolate hypothesis-specific variance.

## Supplementary Materials for

## 1 Formalism for TL and BP

In this section we describe the underlying mathematics of both TL and BP in general terms. Note that practical implementations of either make simplifications or approximations that are useful in practice, which we will not develop in detail.

A learning algorithm for a neuronal network is an optimization procedure that decreases a loss function

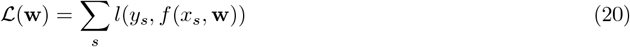

were **w** are the weights of the connections between neurons, *y*_*s*_ is a desired output or activity of the network, and *f*(·,·) is the function implemented by the network parameterized by the weights and the sensory input,and *l*(·,·) is an error function.

Note that both the error function *l* and the loss *L* are mathematical constructs, which can always be defined for any learning process, but do not need to be implemented by any observable biological quantity. The formalism is still useful to make our analysis understandable and to connect our results to the machine learning literature as well as computational neuroscience frameworks such as predictive coding or energy models.

### 1.1 Formalism for BP

For a given sample, we can compute the change of a single weight as

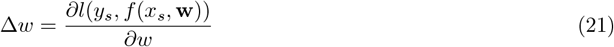

where we can apply the chain rule passing through the activities of the neurons,

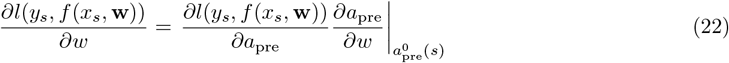

Now we introduce an internal variable in the neuron that linearly reflects the input, *a*_pre_= *ϕ*(*v*_post_), with *ϕ*() being the activation function. This can then be further split into

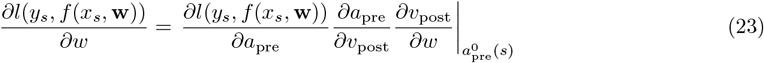

where by construction the internal variable grows linearly with the input, so

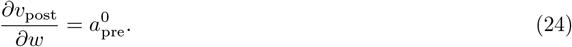

Thus,

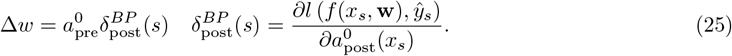

### 1.2 Formalism for TL

In TL the loss is minimized by imposing a different activity through an input to each neuron, so that

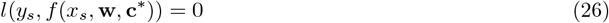

where **c**^*^ is a vector encoding the signal that imposes the activity onto each neuron. Depending on the architecture, this signal is called a control signal, a top-down prediction or a feedback input, and is computed in various ways (Rao and Ballard, 1999; Meulemans et al., 2022a,b; Song et al., 2024; Scellier and Bengio, 2017). In general, the control is chosen so as to provide the smallest input that will fulfill the aforementioned condition. Energy models, for example, follow the direction of steepest descent of the loss (Rao and Ballard, 1999; Scellier and Bengio, 2017), although other solutions such as control theory approaches are also used (Meulemans et al., 2021). In the case of unsupervised learning, *y*_*s*_ is simply the activity of the network that is closest to the one imposed by *x*_*s*_ but fulfills a constraint on the activity such as sparsity.

The main point of TL is to provide a plasticity such that the control input will not be required, and thus the network will have learned the right computation without needing this control.

Now a subtle step is to connect the synaptic weight update to the reduction of the control signal **c**. Since **Eq. 26** is an invariant, the change of loss is zero and so is the input, hence

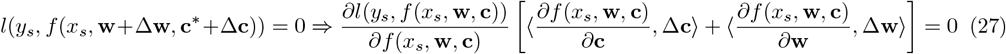

where≈·,·≈ is a dot product, and the derivatives are taken at for a control signal **c**∈ [**c**^*^+ ≈**c**] that comes from the mean value theorem. Assuming that the change in control from one iteration to the next will be small (akin to the assumption in BP that the weight updates impose a small change), **c**≈**c**^*^. From this, we can obtain the constraint

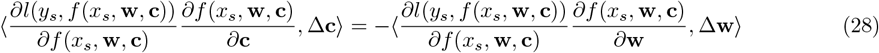

where we can already see how the weight update relates to the control signal, which we will now make more precise and in terms that are similar to the standard machine learning formalism.

The first step is to recognize that both terms in **Eq. 28** are single scalars, but are computed as dot products of high dimensional quantities. Thus, the constraint is on one degree of freedom, but we want to turn the constraint into an optimization process that applies to many neurons and weights.

For this we need to select the weight update that reduces the control signal. Different options are possible, as we can pick any norm on the control or on the activities. More specifically, any weight update where **c** goes to zero, will work. This can be done by minimizing the control or feedback signal Song et al. (2024), the difference between the feedforward and target activities Meulemans et al. (2021), or through temporal hebbian rules Aceituno et al. (2023).

A simple case would be to push the system to the desired activity as fast as possible, in which case

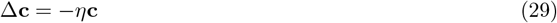

where the proportionality constant corresponds to the learning rate. Thus, we can rephrase **Eq. 28** for single neurons,

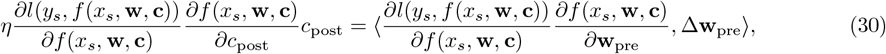

and apply the chain rule to the partial derivatives and make them pass through the postsynaptic activity of the neuron,

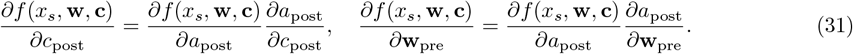

Since there is a derivative that relates the postsynaptic activity to the loss (a scalar to a scalar) on both sides of **Eq. 30**, we can simplify to

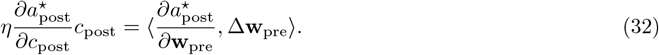

Using again the internal neuron variable that conveys the weighted sum of the neuron inputs,

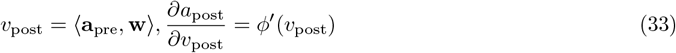

we can then simplify **Eq. 32** to

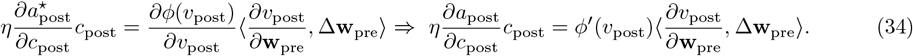

where

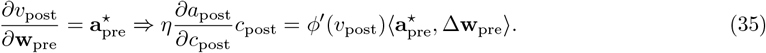

where 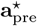 is the target presynaptic activity. Note that a general assumption is that learning has to be done in the full network simultaneously, most TL architectures implicitly use a presynaptic activity that is controlled to be in the target state when learning happens (Rao and Ballard, 1999; Meulemans et al., 2022b; Song et al., 2024).

Just as with the control signal, there are many possible changes of the presynaptic weight that would reduce the amount of control signal for the simple reason that there are usually many presynaptic neurons. Thus, we assume that the learning weight should be efficient, in the sense that the learning should provide the maximum decrease of *c*_post_ for the minimum amount of weight changes Δ**w**_pre_. Geometrically, the natural way to maximize the effects of a presynaptic weight update is to align the update with the presynaptic inputs, thus setting the angle of the vectors in the dot product to zero.

Taken together, the update rule for a single synapse ends up being

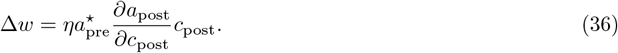

and by using the mean value theorem, we can set

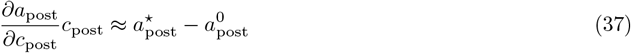

and if we simplify *ϕ*(*v*) to be linear (or a rectified linear unit),

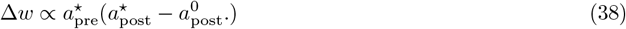

where we then recover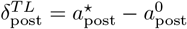

## 2 Links between BP and TL

The two families of algorithms that we study are not separated classes, but rather two extremes of a spectrum. While both algorithms adapt the synaptic weights to modify the activity so that a desired output is provided, BP uses the neuronal responses of the network to a stimulus, while TL disregards the current responses and uses instead the desired neuronal responses. This difference can emerge from the effect of the teaching signal on the neural activity or not (Meulemans et al., 2022b), and indeed various architectures originally designed for TL can be turned into BP in such way (Scellier and Bengio, 2017; Whittington and Bogacz, 2017; Rao and Ballard, 1999).

We make this relationship more precise by showing the possibility to use the machinery of BP to implement TL and vice versa:

- We can use BP to implement TL by using the BP gradients as a control signal (Ernoult et al., 2022). However, this requires adding a parameter to specify the strength of the gradient signal, or multiple iterations of the gradient. In either case, the parameter or iterated gradient needs be re-computed through the learning process, as the nonlinearities in a neural network would imply that the error does not scale linearly with the control signal.
- We can modify TL to approximate BP, by creating ad-hoc targets that are very close to the current activity (LeCun, 1986; Whittington and Bogacz, 2017). However, this requires creating a target for each stimulus presentation, rather than a global target that is maintained through learning (which is how TL is defined).

We can now formalize this statements. The gist of our argument relies on the fact that the transpose of the Jacobian of a system (which is computed in BP) and the inverse of that Jacobian (which provides optimal control) are closely related.

### 2.1 BP with specific strong learning signals is a close approximation of TL

Consider a neural network with a random initialization (as is often done in both deep neural networks and computational models). Through this initialization, we can make the following assumptions:

- All entries of the Jacobian matrix are sampled from a random distribution
- The mean of that random distribution is zero (the network is balanced), and the standard deviation *σ*_pre_ is finite.
- The network is large, and contains many more neurons than outputs.

Our goal is to show that the first order control matrix *Q* and the transpose of the Jacobian are very similar. Starting with the control signal, a control matrix in a linearized system would guarantee that sending every neuron a control signal given by

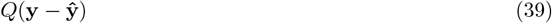

where **y**−**ŷ** are the desired and actual system output respectively. A well conditioned control matrix for that network has the property that

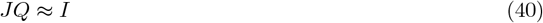

where *I* is the identity matrix (Meulemans et al., 2021), where the rank of *I* is small compared to the number of neurons. Using the transpose of the Jacobian with a scaling as *Q*, we get that the control signal will work if

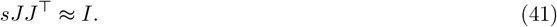

Consider first the product of the Jacobian with its transpose. We can consider the value of entries in the diagonal and off diagonal of the product *P*= *sJJ*^⊤^

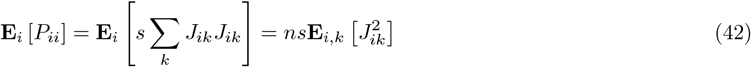

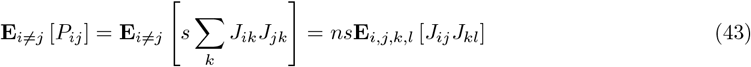

and by using the standard rules to compute the variance and expectation of the product of independent variables,

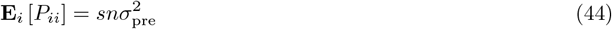

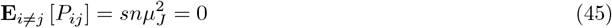

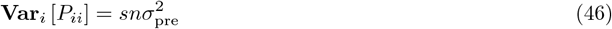

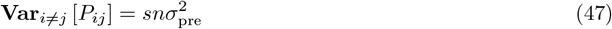

Thus, the matrix *P* is effectively equivalent to

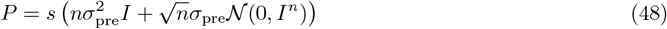

where *N* (0, *I*^*n*^) is a random matrix with gaussian entries. Since 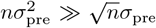, the random component will be small, and we can ensure that *P*≈ *I*, by setting

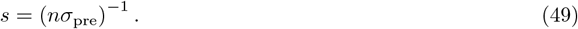

For example, if we set the Jacobian initialization ensuring that *σ*_pre_= *n*^*−*1*/*2^, then *s*= 1.

Thus, for *s*= (*nσ*_pre_)^*−*1^ the gradient signal can be used as a control signal. Note, however, that this requires a specific value of *s*. In other words, for BP to be similar to TL, the two following properties have to be fulfilled:

- The error signal acts as a control signal to the firing of the neuron.
- The error signal is scaled as *s*= (*nσ*_pre_)^*−*1^.

Thus, BP can be interpreted as TL under those assumptions. However, there are two key differences to keep in mind:

- TL is equivalent to BP only if the parameter *s*(the learning signal strength) is set to a precise value, while in TL the value of *s* emerges naturally from first principles. Thus, for such observation TL is more parsimonious and thus the preferred theory.
- The previous assumptions work under linearized dynamics. However, when we have strong nonlinearities the value of *s* would need to be fine-tuned.

#### 2.2 TL with small nudging targets is an approximation of BP

Having shown that BP with strong feedback is close to TL, we now prove the complementary direction and argue that TL with weak feedback is close to BP. For simplicity and to align with our implementation of TL, we will use the nomenclature of control theory, although a similar argument has been build for other specific implementations such as predictive coding (Whittington and Bogacz, 2017) or Equilibrium Propagation (Scellier and Bengio, 2017).

In TL, the learning signal *δ*_*TL*_ from **Eq. 38** is computed as the input necessary to enforce a target activity at the output **ŷ**. Using a Taylor expansion approximation,

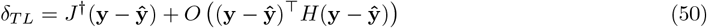

where *J*^*†*^ is a pseudo-inverse of the Jacobian, which accounts for second-order interactions between neurons and *H* a matrix capturing nonlinearities and second-order interactions in the network. Matrix inverses and second order interactions are notably hard to compute, hence to consider the limit **y**− **ŷ**≈ 0 is often more practical. Then the second order components are ignored and

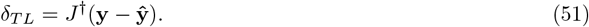

Furthermore, in because of the small error we can simply consider the effect of a small perturbation in the neurons and compute how they would affect the output. This yields the simple computation

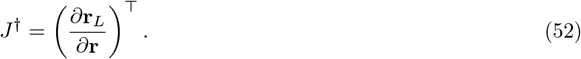

Hence, if we set up a target that is very close to the current output,

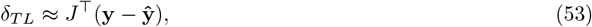

which is the learning signal given by BP.

The natural question is then whether targets that are always closer to the current output are feasible to have. A natural approach is to define intermediate small targets, or “weak nudging” (Meulemans et al., 2021), where the control signal continuously adapt the target to be always close to the current output, but in the direction of the desired output activity.

Using such architectures we can obtain a valid approximation of BP through TL, but then we need to add an extra layer of complexity: on top of the target defined by standard TL, we need to add a circuit that computes a small target that can then be used to compute the learning signal. Thus, considering BP as an approximation of TL is less parsimonious than simply considering BP.

## 3 Derivation of the target learning rule from our PN model

To render our mechanistic PN comparable to the average activities observed in neuronal recordings in vivo, we reformulate our PN model as rate-based with a three-factor learning rule. The inputs that arrive at the soma and the inputs that arrive at the apical dendrites are assumed to follow a Poisson distribution with a relatively low firing rate, in order to ensure that consecutive presynaptic spikes or feedback spikes are temporally separated with high probability, allowing us to consider spikes and bursts as independent events.

We divide the somatic output into bursts and single spikes. The number of events where somatic presynaptic input drove the soma over its firing threshold is represented by *N*. The events will sometimes generate a burst *N*_burst_ and sometimes a single spike *N*_spike_, thus *N*= *N*_burst_+ *N*_spike_. A burst is generated only if feedback input arrives at the apical dendrites within a short time interval Δ*T* after a somatic postsynaptic spike. The aforementioned assumption of low firing rates for apical and somatic inputs, combined with poisson events, implies that somatic and apical spikes rarely arrive very close in time, and thus two events do not lead to the same burst. As a consequence, the probability of turing a single spike into a burst grows linearly with the apical input rate. We can thus calculate the expected number of bursts and single APs as,

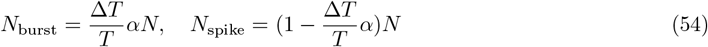

where *T* represents the total time of the simulation, and *α* is the rate of apical input. The number of bursts is simply the fraction of time at which the apical input can generate a burst times the probability of receiving suprathreshold somatic activity.

Lastly, the firing rate of the postsynaptic neuron increases when burst firing occurs. Therefore, considering the activity as the number of spikes, the activity can be expressed as,

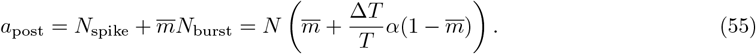

where 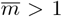 is the expected number of spikes per burst. As denoted here, the expected firing rate of the neuron grows monotonically with the amount of apical feedback.

We can now derive plasticity from the calcium model, understanding that a plateau potential generates LTP, and a burst and a single spike without the corresponding apical input generates LTD when there is a presynaptic spike

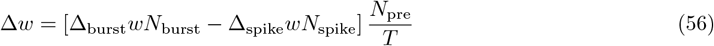

where Δ_burst_*w*> 0 and Δ_spike_*w*> 0 denote the synaptic change per burst or spike, respectively, and *N*_pre_ correspond to the number of presynaptic spikes. Substituting **Eq. 54** into **Eq. 56** produces,

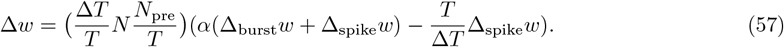

Now there exist a *α*^baseline^ such that Δ*w*= 0, which corresponds to

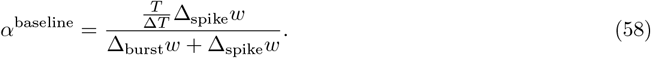

Using *α*^baseline^, we can write the synaptic weight change as

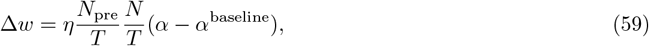

where *η*= Δ*T*(Δ_burst_*w*−Δ_spike_*w*). To simplify the notation further, we will denote the presynaptic firing rate as *r*_pre_= *N*_pre_/*T* and the postsynaptic firing rate at baseline apical input 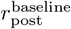, which leaves us with the synaptic update rule

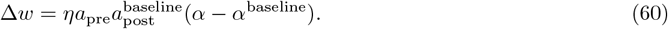

and if we set *α*^baseline^= 1 for simplicity, the postsynaptic firing rate for a given apical input is then

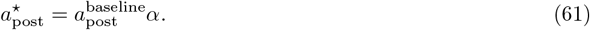

We can then plug this postsynaptic activity into **Eq. 60**, and obtain

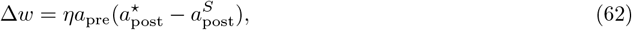

where 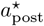 is the target postsynaptic activity which is driven by the apical input. In this formulation the error term matches the one used in existing architectures Rao and Ballard (1999); Whittington and Bogacz (2017); Meulemans et al. (2021), and the learning rule as we tested in **Eq. 5**.

## 4 Apical plasticity

In contrast to our results on basal synapses, we found that for apical synapses neither the plasticity induction AUC (*R*_*Spearman*_= 0.1049, *p*= 0.7456, **Fig. S1D**) nor the number of APs (*R*_*Spearman*_= 0.3996, *p*= 0.1982, **Fig. S1E**) are good predictors for changes in synaptic strength. This difference in the predictability of basal versus apical plasticity is also reflected in the finding, that overall changes in basal EPSP amplitudes did not correlate with changes in apical EPSPs (*R*_*Spearman*_= 0.2308, *p*= 0.4705, **Fig. S1F**).

**Figure S1:**
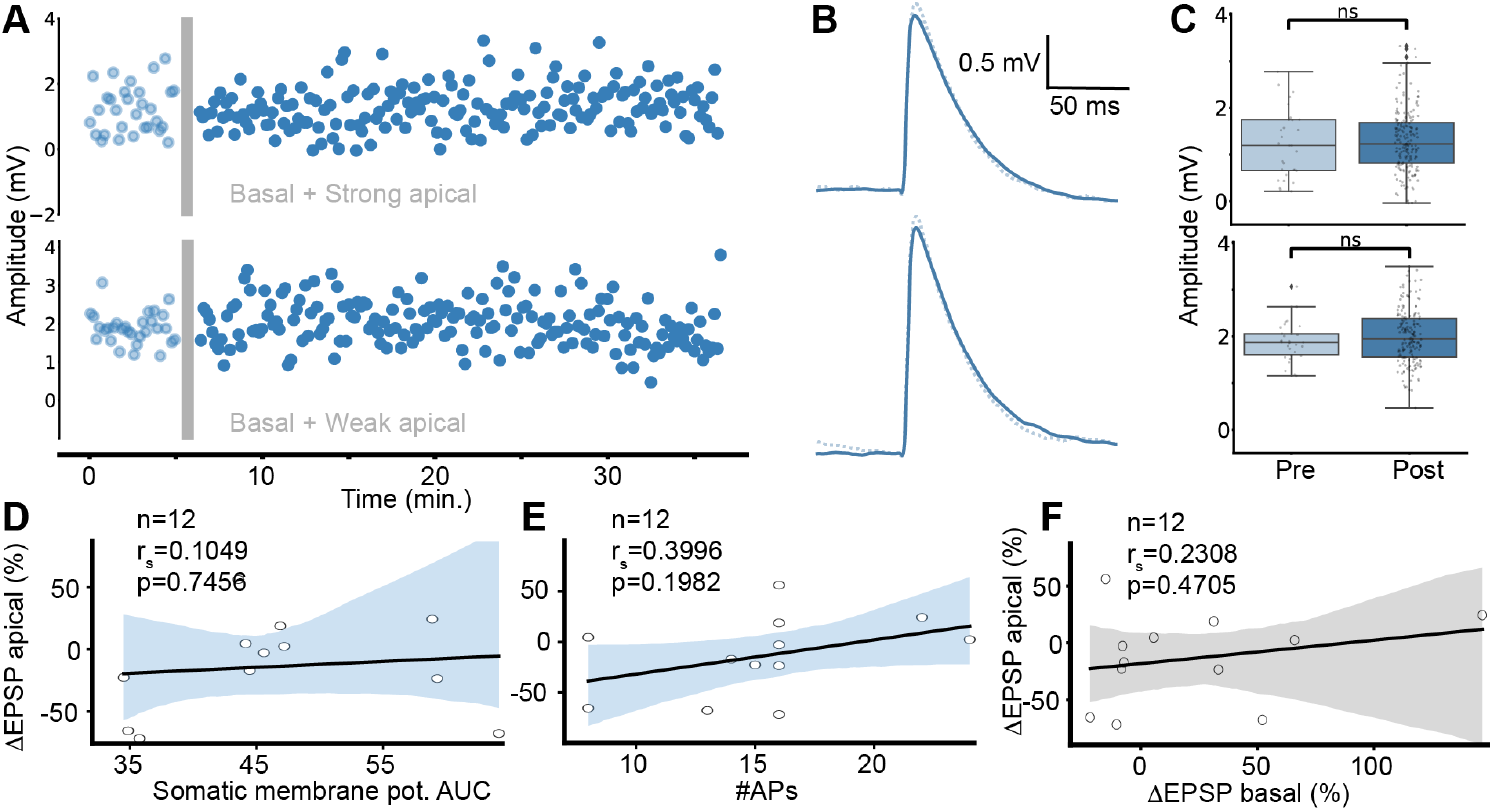
Representative example recordings of EPSP amplitudes during apical stimulation before and after the plasticity induction event. (**A**) EPSP amplitude evolution over time; each dot represents one EPSP pre (light blue) and post (dark blue) plasticity induction event (gray bar). (**B**) The quantification of EPSPs for two sample cells shows no significant change in EPSP amplitudes. (**C, D**) Linear correlation analysis between changes in apical EPSP amplitude and somatic membrane potential AUC ((C) n=12, *r*_*S*_=0.1049, p=0.7456) and number of APs ((D) n=12, *r*_*S*_=0.3996, p=0.1982), respectively. (**E**) Correlation analysis of basal and apical ΔEPSP (n=12, *r*_*S*_=0.3669, *p*=0.2408). (**F**) Correlation between basal and apical EPSP amplitude changes *r*_*s*_= 0.2308, *p*= 0.4705.

**Figure S2:**
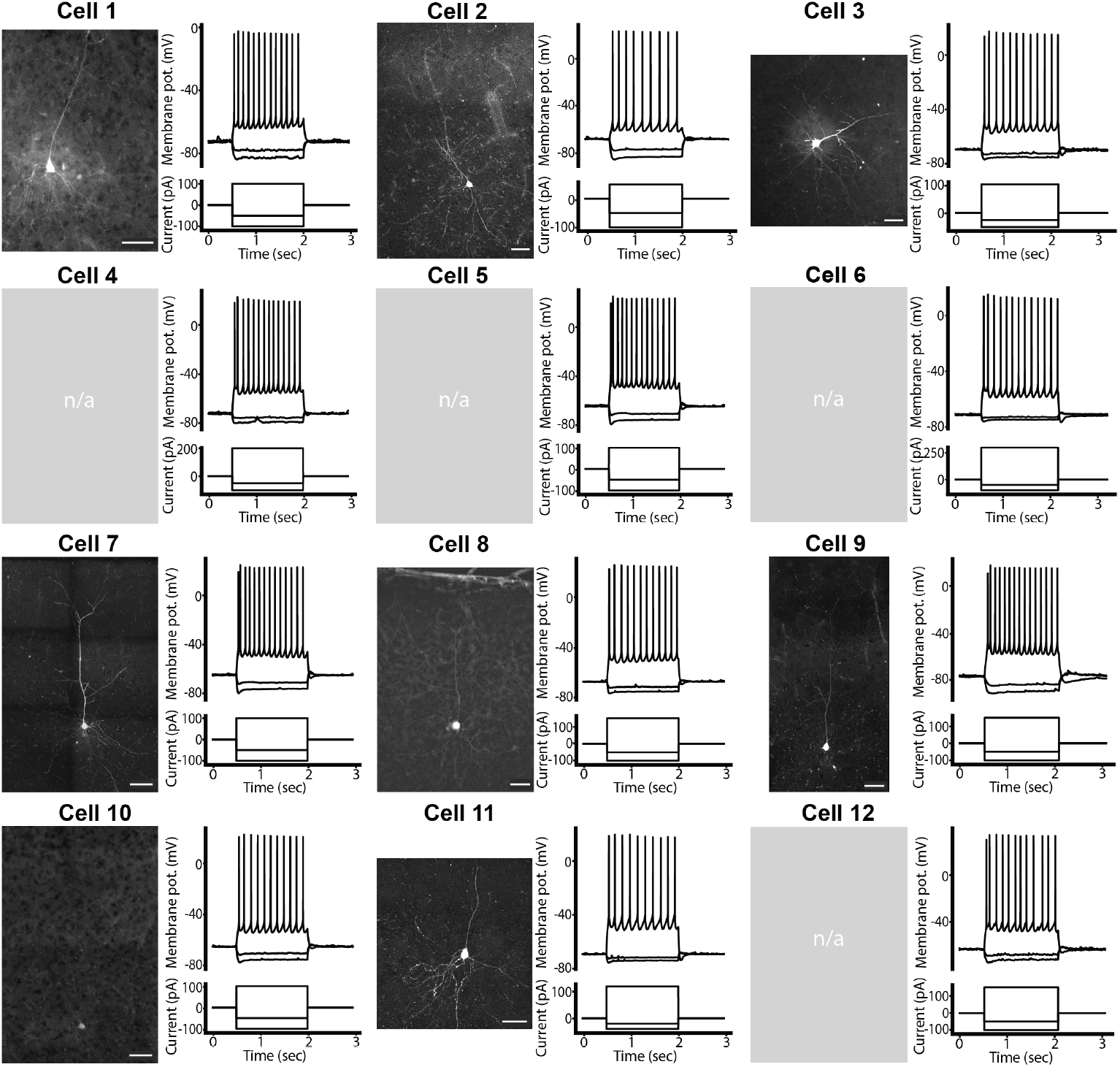
Biocytin-Streptavidin stained neurons (left) and typical hyperpolarizations and action potential trains (right top) induced by negative and positive current injection steps (right bottom) from recorded neurons. For cells 3, 4, 5, and 11, no Biocytin-Streptavidin staining/morphology reconstruction was performed.

**Figure S3:**
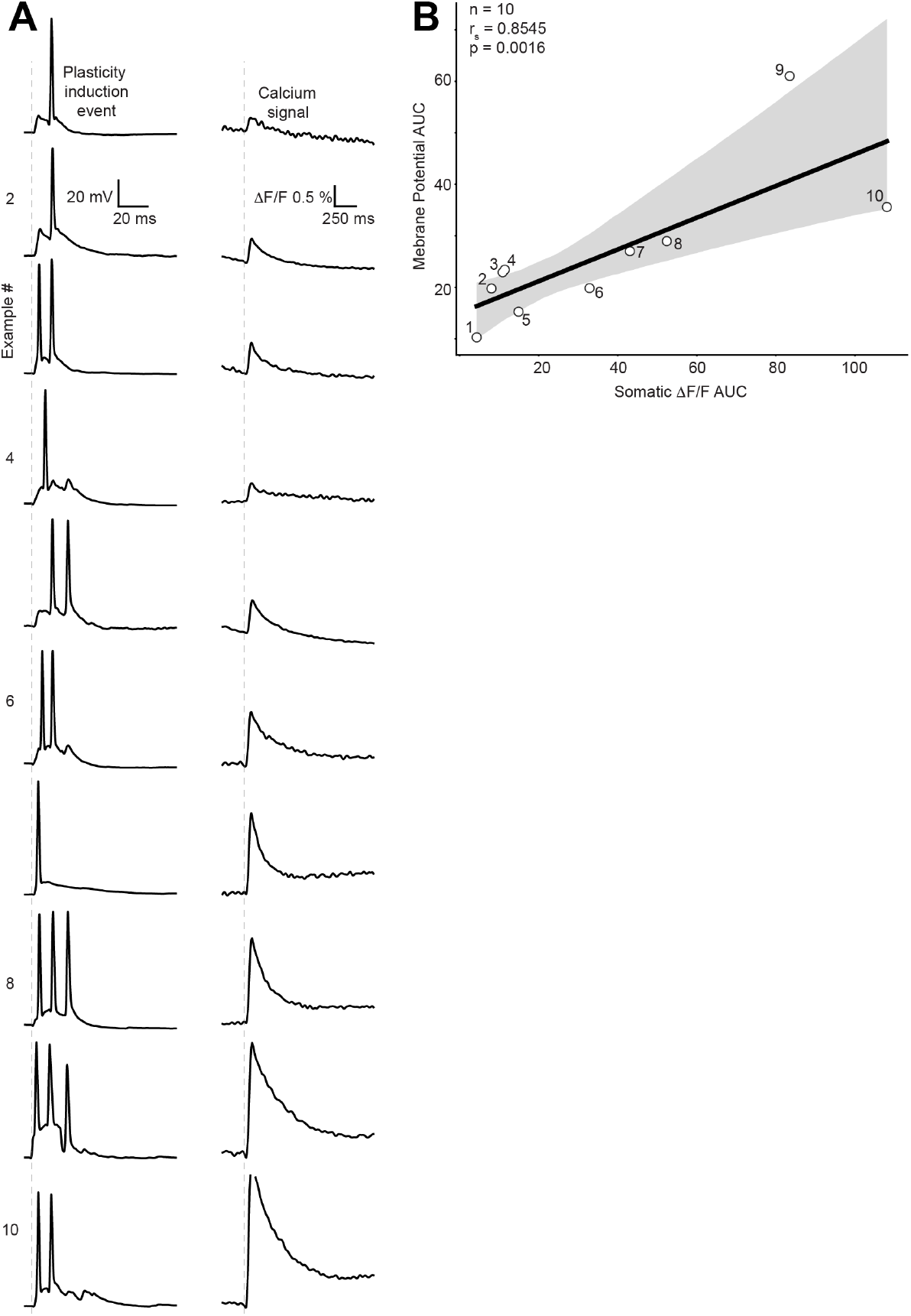
(**A**) Somatic membrane potential recording during plasticity induction (left) and somatic calcium ΔF/F during the same event (right). In total, showing events from 10 recordings in 6 neurons (4 mice) from **Fig. S2**. (**B**) Linear fit of induction event AUCs and somatic calcium signal AUCs showing significant correlation (n=6, 10 events total, *R*_*S*_=0.8303, p=0.0029)

**Figure S4:**
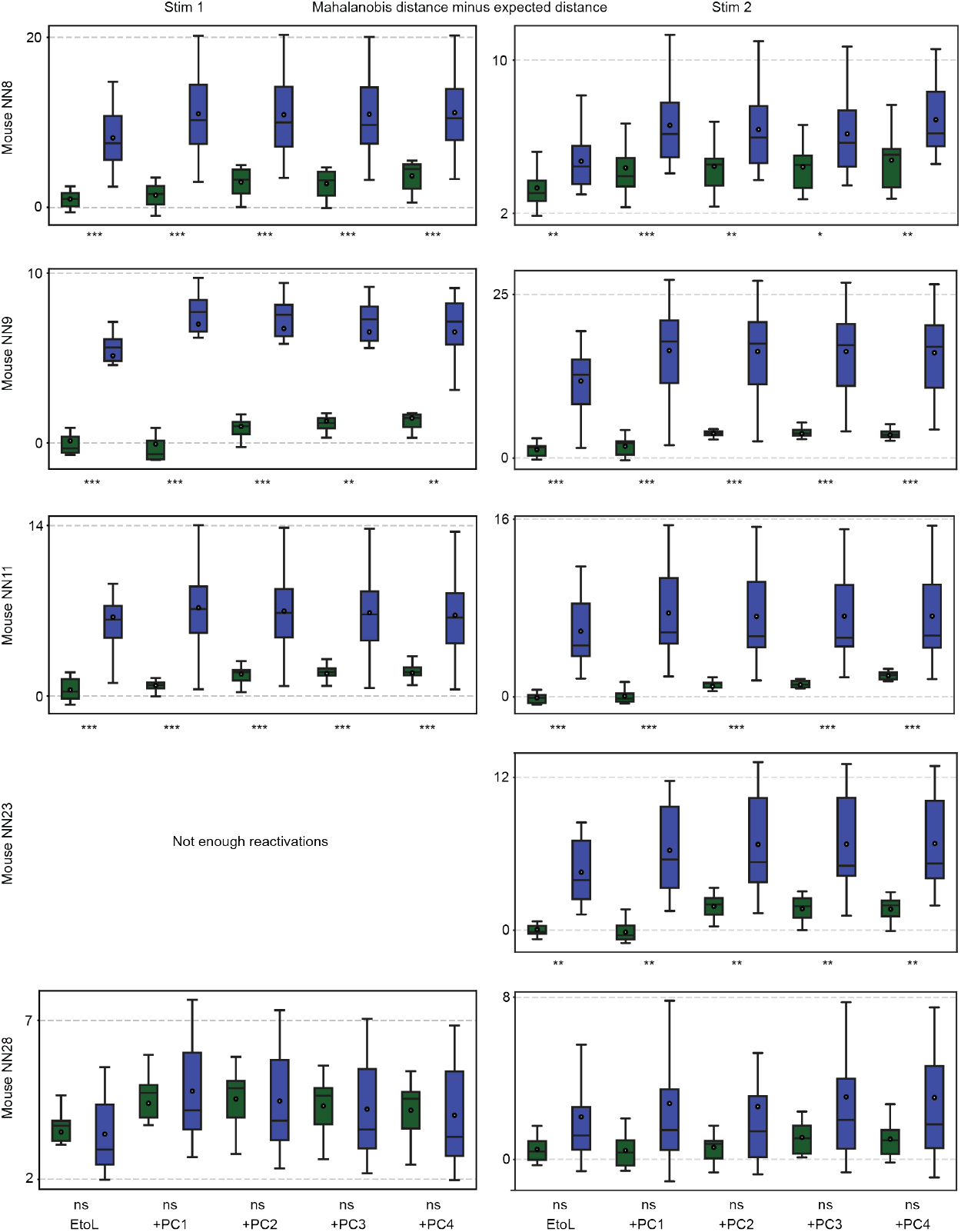
The Mahalanobis distances for the TL (green) and BP (blue) hypotheses for both stimuli and all mice. The statistical significance is denoted beneath the x-axis for each pair of distributions. For each sub-figure, every subsequent pair of distributions denote an additional dimension added to the component space.

